# Differences in adult survival drive divergent demographic responses to a decade of field warming on the Tibetan Plateau

**DOI:** 10.1101/2023.10.03.560467

**Authors:** Hai-Tao Miao, Roberto Salguero-Gómez, Katriona Shea, Joseph A. Keller, Zhenhua Zhang, Jin-Sheng He, Shou-Li Li

## Abstract

1. A central question in biodiversity conservation is whether species will maintain viable population dynamics under future climate change. Assessing species extinction risk under climate warming requires demographic studies integrating vital rate responses to long-term warming throughout species’ life cycle. However, studies of this nature are rare.
2. Here, we examine the demographic responses of two co-occurring herbaceous plants, *Elymus nutans* Griseb. and *Helictotrichon tibeticum* (Roshev.) Holub, after a decade (2011-2020) of *in situ* active warming by 2°C in the grasslands of the Tibetan Plateau. We parameterise Integral Projection Models (IPMs) to project the population dynamics under ambient and long-term warming conditions, and examine the key vital rates responsible for any potential differences in population growth rates.
3. Warming has contrasting effects on the two functionally similar co-occurring species: warming promotes the population growth rate of *H. tibeticum*, but intensifies the population decline of *E. nutans*. Our elasticity analyses show that survival is the most important vital rate for population viability in both species under both ambient and warmed conditions. Furthermore, our retrospective Life Table Response Experiment (LTRE) analysis reveals that the contrasting fates of the two species under warming mainly arise from the different responses of adult survival, which is significantly promoted in *H. tibeticum* but slightly reduced in *E. nutans.* Individual shrinkage occurred 1.6-fold more frequently under warming than ambient conditions for both species, and made considerable negative contributions to their population growth rates in warmed plots. However, such negative effects are offset in *H. tibeticum* (but not *E. nutans*) by the positive contribution to population growth rate of the associated increased survival.
4. *Synthesis*. Our study illustrates that the responses to climate warming may vary considerably between similar co-occurring species, and species with a demographically compensatory strategy may avoid population collapse. Additionally, caution is needed when generalizing findings among functionally similar species, and that conservation measures should be tailored at the species level.

## INTRODUCTION

Climate warming poses a growing threat to global biodiversity by rapidly altering organisms’ environments. Thus, predicting the risks of local extinction, and developing proactive conservation measures to protect species in danger is of increasing urgency (Panetta et al., 2018; Pearson et al., 2014). Species can persist in the face of climate warming if they are able to maintain viable populations under novel climate conditions (Nomoto & Alexander, 2021; Peterson et al., 2018). However, quantitively assessing the extent to which climate warming can shape population viability is challenging, as it requires demographic studies integrating vital rates (*e.g.*, survival, growth, and reproduction) over an entire life cycle in response to long-term warming. The impacts of climate warming on individual performance in terms of changes in morphology, physiology, phenology, and vital rates have been extensively studied (Blumenthal et al., 2016; Gornish et al., 2014; Wang et al., 2020). However, we still lack mechanistic insights into the impacts of such changes on long-term population viability (Iler et al., 2021; Selwood et al., 2015). This knowledge gap hinders our capacity to strategically manage populations of conservation concern under future climate change (DeMarche et al., 2018; Nomoto & Alexander, 2021; Keller & Shea, 2021).

The direct effects of climate warming on individual performance do not directly scale up into impacts on population viability (Iler et al., 2021; McLean et al., 2016; Selwood et al., 2015). Climate warming may affect vital rates differently, not only in magnitudes but also in direction (Keller & Shea, 2021; Nomoto & Alexander, 2021; Panetta et al., 2018). When the negative effects of climate warming on some vital rates were compensated by positive effects on others, then the overall effects at the population level could be alleviated or cancelled out, a phenomenon called “demographic compensation” (Villellas et al., 2015; Sheth & Angert, 2018). As such, evaluating the impacts of climate warming on species viability requires integrating individual- and population-level responses. The presence of demographic compensation among vital rates may enable populations to persist under climate warming. For example, a recent study on the alpine plant *Silene acaulis* found that the reduction in survival was compensated by the increase in growth under increased temperature toward lower latitudes (Peterson et al., 2018). Such demographic compensation allowed *S. acaulis* to maintain viable populations across its range (Peterson et al., 2018). Alternatively, populations lacking demographic compensation are more likely to decline under elevated temperatures (Cui et al., 2018). Additionally, the extent to which population persistence is affected by climate change is determined not only by the magnitude of warming-induced changes in vital rates, but also by the sensitivity of population growth to such changes. This relationship is well known not to be linear (de Kroon et al., 2000; Villellas et al., 2015). As such, assessments of population viability require more comprehensive modelling efforts to assess the cumulative effects.

Another challenge hampering investigation of demographic impacts of climate warming arises because such evaluations require demographic studies under long-term warming. It may take multiple years for populations, especially populations of perennials, to respond to elevated temperature (Evers et al., 2021; Tenhumberg et al., 2018). Existing findings about the impacts of climate warming on population viability are inconsistent, with negative and positive effects being widely reported at the population level in different systems (Gornish et al., 2014; Keller & Shea, 2021; Nomoto & Alexander, 2021; Panetta et al., 2018; Reed et al., 2021). For example, Keller and Shea (2021) found that the population growth of the globally invasive plant *Carduus nutans* was greatly accelerated under a short-term warming treatment; a study by Gornish et al. (2014) showed that short-term warming increased the growth of the populations of pineland silkgrass *Pityopsis aspera*; and Panetta et al. (2018) reported that a 25-year warming treatment caused declines in the population size of a widespread mountain plant *Androsace septentrionalis* L.. One reason for such inconsistent findings may be that most results are based on short-term warming experiments (but see Panetta et al., 2018), often shorter than the lifespan of the study species. These treatments may only capture the transient dynamics of the populations under warming, while plant populations may respond over the span of multiple years (Evers et al., 2021; Zhang et al., 2012), or sometimes even decades (Panetta et al., 2018; Tenhumberg et al., 2018).

Alpine grasslands are considered among the most vulnerable ecosystems to climate warming (IPCC, 2019; Verrall & Pickering, 2020). In particular, the annual temperature on the Tibetan Plateau has increased at a rate twice the global average in the past 50 years (Chen et al., 2013). Therefore, exploring the consequences of climate warming on population persistence on the Tibetan Plateau is important to inform the management of such critical susceptible alpine ecosystems. Moreover, such findings can serve as early warning signals for the conservation of other ecosystems. Previous studies on the grasslands of the Tibetan plateau have found that long-term warming advanced phenology and shifted plant community composition toward an increased abundance of herbaceous functional groups (Liu et al., 2018; Wang et al., 2020). However, it remains unknown how the maintenance of alpine plant populations on the Tibetan Plateau will be affected by climate warming and whether functionally similar species inhabiting the same habitats would show similar demographic responses to elevated temperatures.

Here, we examine how climate warming may affect the population dynamics of two dominant plant species, *Elymus nutans* Griseb. and *Helictotrichon tibeticum* (Roshev.) Holub, on the alpine grassland of the Tibetan Plateau. Our study sites have received a 2℃ active warming starting in 2011. To quantify the responses of both species to warming, we parameterised Integral Projection Models (IPMs; Easterling et al., 2000) with demographic data collected in 2019 and 2020. We used these models to test the following hypotheses: i) Long-term warming may negatively affect the individual vital rates and subsequently reduce the long-term population growth rate, because alpine plants are generally vulnerable to elevated temperatures; ii) The species with demographic compensation among vital rates may alleviate the warming-induced negative effects on population growth and thus save populations from collapse; iii) The co-occurring species may adopt different demographic strategies to cope with elevated temperature; if so, conservation measures should be tailored at species level under future climate warming.

## MATERIALS AND METHODS

### Study area and experimental design

The study was conducted at the Haibei National Alpine Grassland Ecosystem Research Station, Qinghai Province, China (37°36′ N, 101°19′ E, 3,125 m a.s.l.). The study area is located on the northeastern Tibetan Plateau. The climate is characterized by a long, cold winter and a short, cool summer. The mean annual temperature is -1.1 ℃, with the highest monthly temperature occurring in July (10.4 ℃) and the lowest monthly temperature in January (-14.6 ℃). The mean annual precipitation is 488 mm, with >80% occurring as rain during growing season, from May to September (Wang et al., 2020). The site is a typical alpine grassland, and the vegetation is dominated by perennial herbaceous plants, such as *Elymus nutans*, *Helictotrichon tibeticum*, and *Stipa aliena* (Liu et al., 2018). The study site was fenced annually in summer (from June to September) to avoid animal disturbance, but was grazed annually in winter by sheep (from October to May) to mimic historic grazing regimes in the region (Fuller et al., 2017).

A warming manipulation experiment was set up in July 2011. Two warming treatment plots and two ambient plots, each of 2.2 m × 1.8 m, were randomly distributed on the landscape with a spacing of about 10 m. The temperature at the warming treatment plots was elevated by 2℃ via two infrared heaters (220 V, 1,200 W; 1.0 m long, 0.22 m wide), suspended in parallel at 1.6 m above the ground. Dummy heaters with the same shape and size as the infrared heaters were used to mimic the shading effect in the ambient plots.

### Study species and demographic data

To compare the effects of climate warming on the population performance of functionally similar plant species inhabiting the same habitats, we selected two co-occurring alpine species, *Elymus nutans* and *Helictotrichon tibeticum*. These two graminoid species have erect culms, but differ in leaf blades and spikes. Both species can reach a height of 1 m at the fruiting stage, but *H. tibeticum* often grows taller and more densely tufted than *E. nutans*. The two species have similar life histories: both are perennial and sprout new leaves annually, regrow in April-May, flower in June-July, and set fruit in August-September. Recruitment is generally from seed, with germination occurring in spring. Sexual maturity is typically attained in the second year after germination. The two species are well adapted to the cold alpine climate, are widely distributed on the grasslands of the Tibetan Plateau, and serve as important forage species (Wang et al., 2020).

To examine the effect of elevated temperature on demographic processes, annual censuses were conducted in August 2019 and 2020 for both species. In the first census, we measured individual plant height, typically a good predictor of vital rates in perennial grasses (Struckman et al., 2019). We also recorded the reproductive status of each individual. To estimate seed production of *E. nutans*, we first measured spike length for each individual, and then randomly selected 50 individuals close (<10 m) to our experimental site and correlated the number of seeds (destructive count) to spike length. Because *H. tibeticum* can produce over 10 spikes per individual, we only counted the number of spikes and later related the number of new seedlings in time *t* + 1 to the number of spikes in *t* at each plot, as oftentimes done in plant demographic studies (*e.g.*, Lucas et al., 2008). Upon the first measurement, we tagged each individual with a stainless steel label and recorded its Cartesian coordinates within the plots to allow tracking through time. In 2020, we checked the survival of all tagged individuals and re-measured size and the aforementioned reproductive characteristics on surviving ones. At that point too, we also located and measured new recruits within our plots.

### Statistical analyses

To examine the effects of warming treatment and initial plant size on vital rates, we employed multiple regression models to relate survival, size changes (growth/shrinkage), and reproduction to the current height and warming treatment. The probabilities of survival *s*(*z*) and flowering *p*_*f*_(*z*) were modelled as logistic regressions, with individual height (*z*) and warming treatment as independent variables. Size changes *g*(*z*^′^, *z*) were modelled using multiple linear regression with future height (*z*′) as response variable against current height (*z*) and warming treatment. Subsequently, variances from the size change regression were again related to current height (*z*) and treatment in another multiple linear regression to test for assumptions of constancy as a function of height on variance. The estimated number of seeds in *E. nutans* or the number of spikes in *H. tibeticum f*_*n*_(*z*) were modelled using Poisson regressions, with current height (*z*) and treatment as independent variables. We performed Wald chi-square tests to examine the fixed effects of current height and warming treatment in the regressions of all vital rates (Fox, 2016).

Size shrinkage, an important demographic process (Salguero-Gómez & Casper, 2010), was observed in both species. To examine the effect of climate warming on size shrinkage, we modelled the proportional occurrence of shrinkage for these shrunken individuals employing a generalized linear model using the beta family with a logit-link function, where warming treatment was included as an independent variable (Cribari-Neto & Zeileis, 2010). We then applied a Wald chi-square test to examine the significance of warming treatment in the model (Fox, 2016). Furthermore, we calculated the intensity of shrinkage for these shrunken individuals by subtracting the individual height in 2020 from that in 2019. Because the data of height shrinkage did not meet the normality assumption, we performed a Wilcoxon signed rank test to examine the effects of warming on the intensity of shrinkage (Wilcoxon, 1945).

### Integral projection models

To examine the overall impacts of climate warming on the population dynamics of *E. nutans* and *H. tibeticum*, we built Integral Projection Models (IPMs; Easterling et al., 2000). IPMs are stage-structured population models where the population is organized according to one or more continuous traits (here height) that may change in discrete time (Ellner et al., 2016). In an IPM, vital rates are included as regression functions of the predicting trait(s), as detailed in the section above. For our purposes, variation in vital rates between ambient and warming treatments can be easily included in the corresponding regression models (Nomoto & Alexander, 2021). In the IPM, the population dynamics of our two species are then mathematically described as:

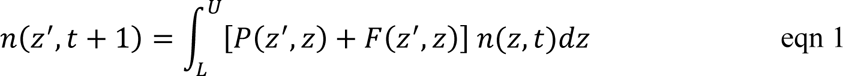

where [*L*, *U*] is the range of all possible heights (*L*=4.6 cm, *U*=102.2 cm for *E. nutans*; *L*=12.9 cm, *U*=124.9 cm for *H. tibeticum*), *n*(*z*, *t*) is the distribution of height *z* at time *t*, and *n*(*z*′, *t* + 1) is the distribution of height *z*′ at time *t* + 1. The expression *P*(*z*^′^, *z*) + *F*(*z*^′^, *z*) is also referred to as the kernel *K*(*z*^′^, *z*), a non-negative surface describing all possible demographic transitions from height *z* at time *t* to height *z*′ at time *t*+1. The sub-kernel *P*(*z*^′^, *z*) comprises all possible transitions of surviving individuals from height *z* at time *t* to height *z*′ at time *t* + 1. This sub-kernel can be decomposed into two functions that determine the probability of survival of an individual at height *z*, *s*(*z*), and the likelihood that the individual will grow/shrink from height *z* to height *z*′ over a year, such that *P*(*z*^′^, *z*) = *s*(*z*) *g*(*z*^′^, *z*). The sub-kernel *F*(*z*^′^, *z*) comprises all sexual contributions of reproductive individuals in time *t* to established recruits in *t* + 1. Here, this sub-kernel is decomposed into four functions including the probability of reproduction *p*_*f*_(*z*), the number of seeds or spikes *f*_*n*_(*z*), the probability a recruit establishes *p*_*e*_, and the probability distribution of recruit size *f*_*d*_(*z*^′^), such that *F*(*z*^′^, *z*) = *p*_*f*_(*z*) *f*_*n*_(*z*) *p*_*e*_ *f*_*d*_(*z*^′^). Because warming significantly affected the vital rate functions (Table S1), we built separate kernels for populations under warmed and ambient conditions for both species.

To examine the effects of climate warming on long-term population viability, we calculated the population growth rate (*λ*) of both species under ambient and warmed conditions. To do so, first, we discretized the IPM into 100 × 100 matrices using midpoint rule (Ellner et al., 2016). The population growth rate calculated from these matrices, their dominant eigenvalue (Caswell, 2001), informs on whether a population will increase (*λ* > 1) or decline to extinction (*λ* < 1) under conditions of stationary equilibrium. To estimate the uncertainty around *λ*, we bootstrapped the individual data 1,000 times to obtain bias-corrected 95% confidence intervals under ambient and warmed conditions for the two study species (Efron, 1987; Anderson & Wadgymar, 2020). Furthermore, we calculated the differences between *λ* values (*Δλ*) estimated in each bootstrap to obtain bias-corrected 95% confidence interval of *Δλ* between ambient and warmed conditions for both species. In addition, to examine whether the demographic responses to climate warming may be affected by a plant’s lifespan, we calculated the mean life expectancy under warmed and ambient conditions for both species, using the *Rage* package (Jones et al., 2022) in R.

To examine the relative importance of each vital rate on the population growth rate to guide population management under climate warming, we conducted elasticity analyses. Elasticity analysis of vital rate functions describes how much *λ* may change when the underlying vital rate changes by a given fraction near a particular size (Ellner et al., 2016; Griffith, 2017; Zuidema & Franco, 2001). By conducting vital rate elasticity analyses, we were able to determine the elasticity values for survival, growth and reproduction separately (Griffith, 2017; Li et al., 2011). Positive elasticity values imply that an increase in a vital rate will increase *λ,* which is particularly useful for distinguishing the relative importance of positive growth *versus* shrinkage. To test how climate warming-induced changes in vital rates might have led to differences in *λ* under ambient and warming conditions, we conducted retrospective demographic analyses, the Life Table Response Experiment (LTRE, hereafter). An LTRE analysis decomposes the observed difference in population growth rate into the contributions from different vital rates (Caswell, 1989). We conducted a fixed-design LTRE on vital rates over a life cycle. The one-factor LTRE analysis follows the formulation:

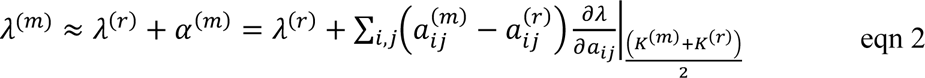

where *λ*^(*m*)^ is the population growth rate under warming treatment *m*, and *λ*^(*r*)^, as a reference, is the population growth rate under ambient conditions. The factor *α*^(*m*)^evaluates the effect of warming treatment *m* on population maintenance. The contribution of different vital rates is calculated by the differences between the vital rate 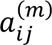 of the large transition matrix *K*^(*m*)^ under warming treatment and the vital rate 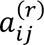 of the large transition matrix *K*^(*r*)^ under ambient conditions multiplied by the sensitivity values of the mid matrix, which is the average of the matrices under warming and ambient conditions (Caswell, 1989). The evaluation of the mid matrix between *m* and *r* minimizes deviances due to the non-linear nature of vital rate sensitivities (Caswell, 1989).

All analyses were performed with the software R 4.0.5 (R Core Team, 2021).

## RESULTS

### The impacts of warming on survival, growth and reproduction

To examine the effects of climate warming on the demographic processes of two co-occurring species, *Elymus nutans* and *Helictotrichon tibeticum,* we used regression models to relate vital rates to current height and warming treatment. We found that the impacts of warming differed among vital rates, across plant sizes, and between species (Table 1). The response of survival to warming was different between *E. nutans* and *H. tibeticum* (Figure 1a,b). Warming increased the survival of shorter individuals but decreased that of taller individuals in *E. nutans* (Figure 1a), whereas warming consistently increased the survival of *H. tibeticum* across all sizes (Figure 1b). The survival of *E. nutans* increased with plant height under ambient conditions but decreased with plant height under warmed conditions (Figure 1a), while the survival of *H. tibeticum* increased with plant height under both warmed and ambient conditions (Figure 1b).

**Figure 1.**
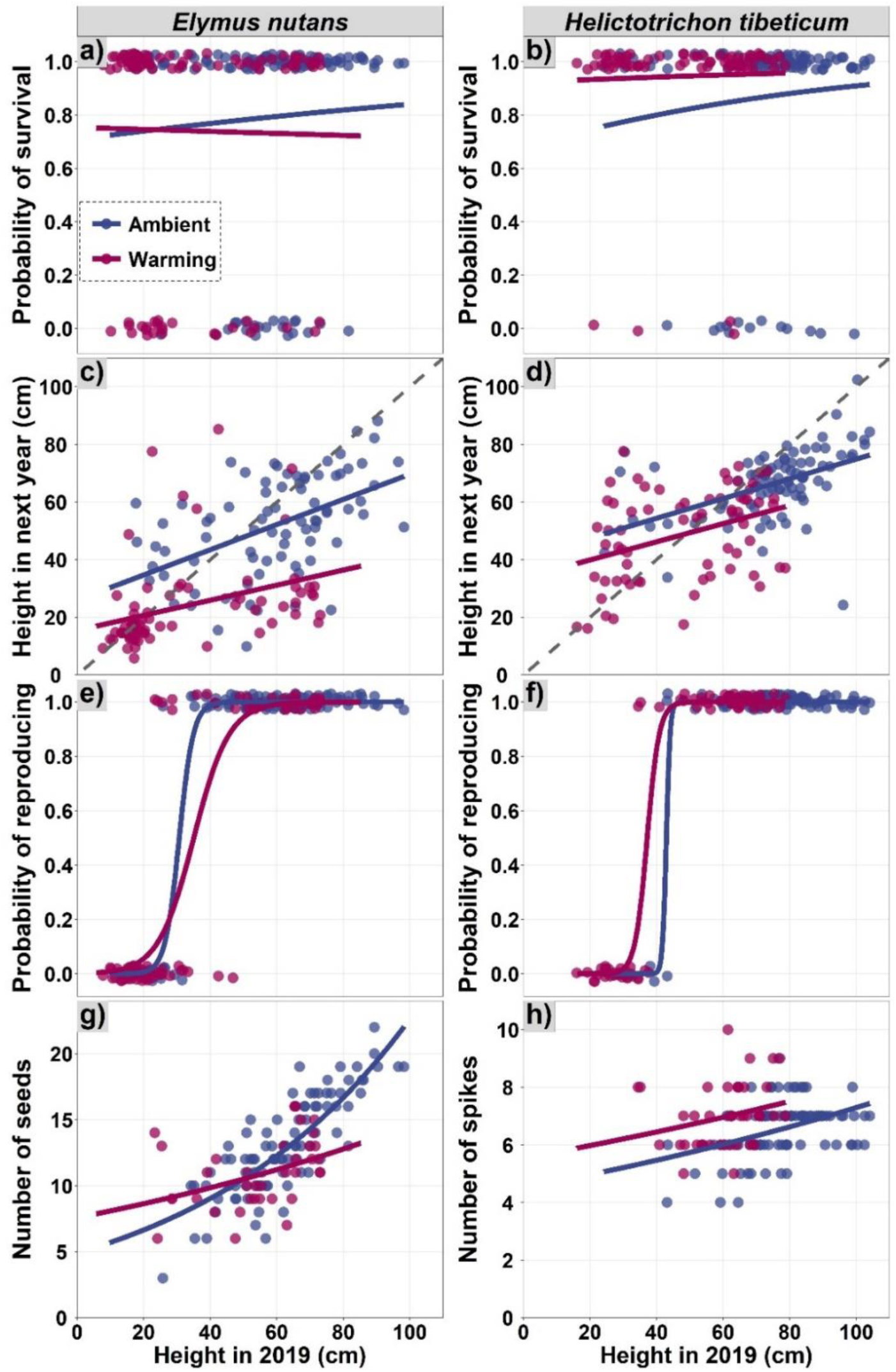
The effects of a decade of warming treatment on the relationships between the vital rates of survival (a, b), growth (c, d), reproduction (e, f), and number of seeds/spikes per capita (g, h), and plant height for two co-occurring alpine herbaceous species, *Elymus nutans* (a, c, e, g) and *Helictotrichon tibeticum* (b, d, f, h), on the Tibetan Plateau. The dashed grey lines in (c) and (d) represent 1:1 line. Regression functions and parameters are given in Table 1.

**Table 1.** Statistical models and parameter estimates used to construct the kernels of the integral projection models for two co-occurring alpine herbaceous species, *Elymus nutans* and *Helictotrichon tibeticum* under ambient and warmed conditions on the Tibetan Plateau.

| Species | Demographic process | Model |
| --- | --- | --- |
| *Elymus*<br>*nutans* | Survival probability ($s$) | Ambient: $\text{Logit}(s) = 0.890 + 0.008z$<br>Warming: $\text{Logit}(s) = 1.115 - 0.002z$<br>$R^2 = 0.005, P < 0.001$ |
| | Changes in height ($g$) | Ambient: $g(z') = 25.967 + 0.437z$<br>Warming: $g(z') = 15.435 + 0.262z$<br>$R^2 = 0.527, P < 0.001$ |
| | Probability of reproducing ($p_b$) | Ambient: $\text{Logit}(p_b) = -14.402 + 0.469z$<br>Warming: $\text{Logit}(p_b) = -6.581 + 0.188z$<br>$R^2 = 0.843, P < 0.001$ |
| | Number of seeds ($b$) | Ambient: $\log(b) = 1.589 + 0.015z$<br>Warming: $\log(b) = 2.026 + 0.007z$<br>$R^2 = 0.709, P < 0.001$ |
| *Helictotrichon*<br>*tibeticum* | Survival probability ($s$) | Ambient: $\text{Logit}(s) = 0.774 + 0.015z$<br>Warming: $\text{Logit}(s) = 2.468 + 0.008z$<br>$R^2 = 0.021, P < 0.001$ |
| | Changes in height ($g$) | Ambient: $g(z') = 40.672 + 0.343z$<br>Warming: $g(z') = 33.618 + 0.315z$<br>$R^2 = 0.414, P < 0.001$ |
| | Probability of reproducing ($p_b$) | Ambient: $\text{Logit}(p_b) = -74.631 + 1.738z$<br>Warming: $\text{Logit}(p_b) = -20.439 + 0.009z$<br>$R^2 = 0.917, P < 0.001$ |
| | Number of spikes ($b$) | Ambient: $\text{Log}(b) = 1.507 + 0.005z$<br>Warming: $\text{Log}(b) = 1.712 + 0.004z$<br>$R^2 = 0.176, P < 0.001$ |

Warming had a negative effect on growth in both study species (Figure 1c,d). In general, changes in height shifted from growth to shrinkage in taller plants (as indicated by the fitted regression lines falling below the 1:1 line for large plants in Figure 1c,d). The frequency of shrinkage was higher under warmed than under ambient conditions, especially for reproductive individuals (Table S3; Figure S1).

The probability of reproducing generally increased with plant height in both species (Figure 1e,f). However, warming increased the probability of reproducing for short individuals in both species (Table 1; Figure 1e,f). Warming increased the threshold of height at which individuals reached 50% probability of reproducing for *E. nutans*, but decreased that threshold for *H. tibeticum* (Figure 1e,f). Warming increased the seed production of shorter individuals but decreased that of taller individuals in *E. nutans* (Figure 1g). However, warming consistently increased the number of spikes in *H. tibeticum* (Figure 1h). Additionally, warming strongly reduced seeding recruitment in *E. nutans* (Figure 2c,g), but not in *H. tibeticum* (Figure 2d,h).

**Figure 2.**
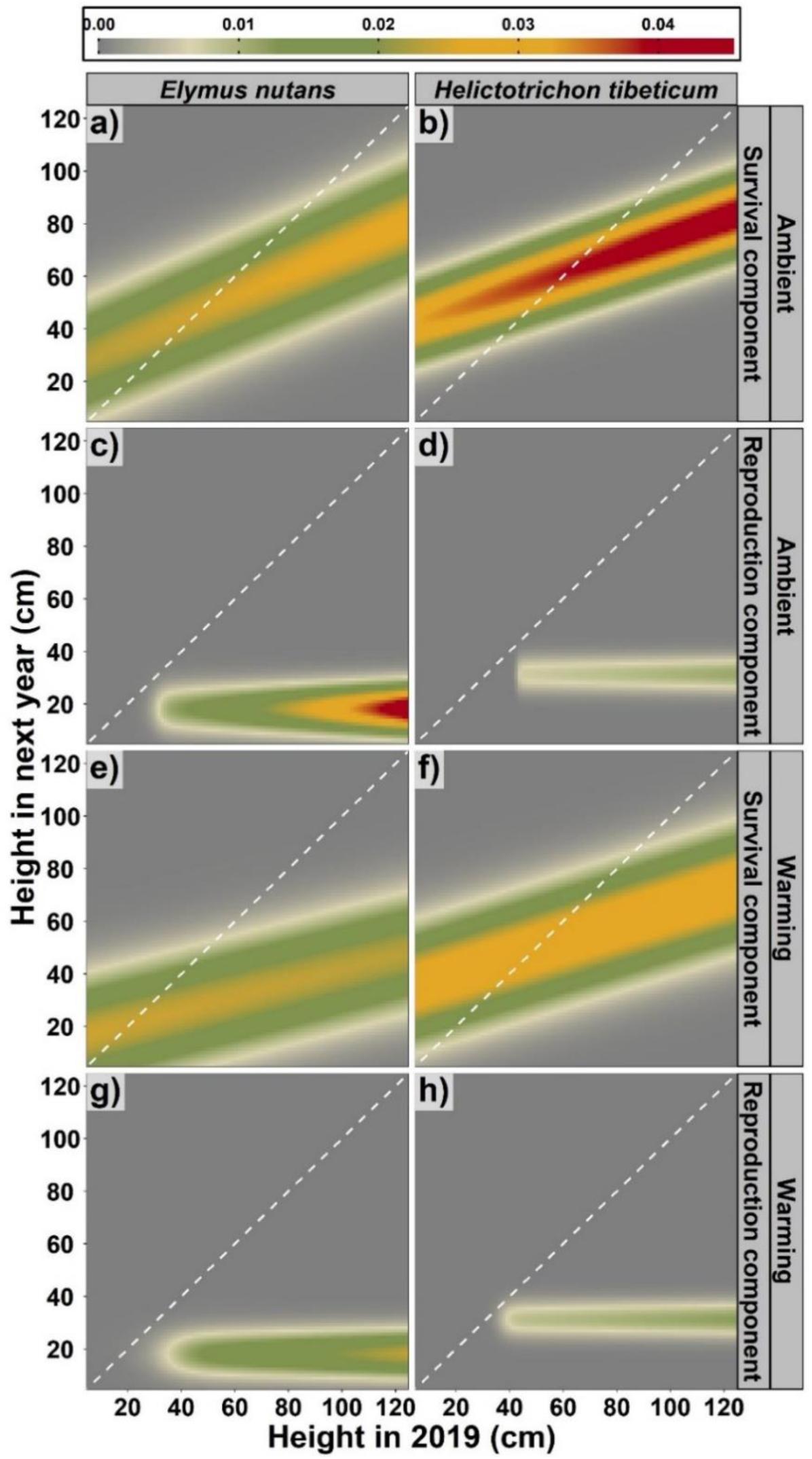
The effects of a decade of warming treatment on the survival components (a, b, e, f) and reproduction components (c, d, g, h) of the fitted kernels of integral projection models for two co-occurring alpine herbaceous species, *Elymus nutans* (a, c, e, g) and *Helictotrichon tibeticum* (b, d, f, h), on the Tibetan Plateau. The dashed 1:1 line along the diagonal represent individuals during the first census that remained the same height or produced seedlings of the same height during the second census.

### The impacts of warming on population growth rate and life expectancy

To examine the effects of elevated temperature on population maintenance, we built Integral Projection Models (IPMs) to estimate the population growth rate (*λ*) for *E. nutans* and *H. tibeticum*. We found that, under ambient conditions, *λ* values were similar for *E. nutans* (*λ* = 0.883, 95% CI [0.879, 0.886]; Table 2) and *H. tibeticum* (*λ* = 0.910, 95% CI [0.907, 0.973]; Table 2). However, the population growth rate of the two species were significantly different under warming: warming decreased the population growth rate of *E. nutans* to 0.694 (*Δλ* = -0.189, 95% CI [-0.193, -0.184]; Table 2), but promoted the population growth rate of *H. tibeticum* to 1.001 (*Δλ* = 0.091, 95% CI [0.087, 0.094]; Table 2). The life expectancy estimated by IPM was similar for the two species under ambient conditions: 4.5 years in *E. nutans* and 6.7 years in *H. tibeticum* (Table S2). However, warming decreased the life expectancy of *E. nutans* to 3.0 years, but increased that of *H. tibeticum* to 15.4 years (Table S2).

**Table 2.** The effects of a decade of warming treatment on the population growth rate (*λ*), difference between *λ* values (*Δλ*) under ambient and warmed conditions for two co-occurring alpine herbaceous species, *Elymus nutans* and *Helictotrichon tibeticum*, on the Tibetan Plateau. Values in brackets refer to 95% confidence intervals (CI).

| Species | Treatments | $\lambda$ | $\Delta\lambda$ |
| --- | --- | --- | --- |
| *E. nutans* | Ambient | 0.883 [0.879, 0.886] | -0.189 [-0.193, - 0.184] |
|  | Warming | 0.694 [0.692, 0.697] |  |
| *H. tibeticum* | Ambient | 0.910 [0.907, 0.973] | 0.091 [0.087, 0.094] |
|  | Warming | 1.001 [0.999, 1.002] |  |

### Vital rate elasticity analyses

To identify which vital rates are key to long-term population maintenance, we conducted elasticity analyses. We found that the population growth rate was overwhelmingly most elastic to changes in survival in both species, under both warmed and ambient conditions (Figure 3a-d). Under warmed conditions, the elasticity to changes in survival increased in shorter plants and decreased in taller individuals in both species (Figure 3c,d). The elasticity to changes in height (*i.e*., positive growth) and probability of reproduction was positive but much smaller than that of survival in both species under both warmed and ambient conditions (Figure 3a-d). The elasticity to shrinkage was negative in all populations, indicating a negative effect on population maintenance as a result of individuals becoming shorter (Figure 3a-d).

**Figure 3.**
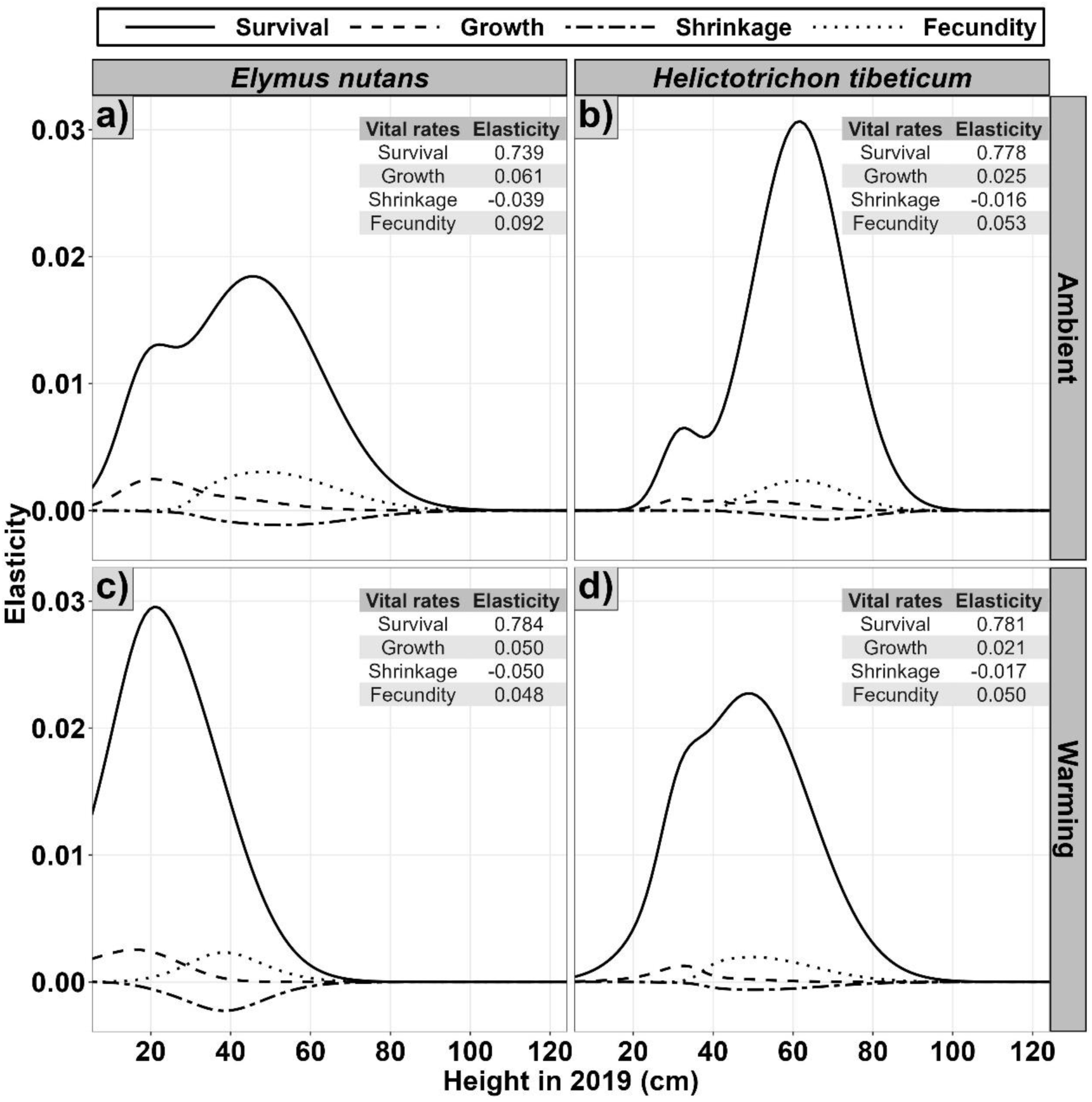
The effects of a decade of warming treatment on the vital rate elasticity of two co-occurring alpine herbaceous species, *Elymus nutans* (a, c) and *Helictotrichon tibeticum* (b, d), on the Tibetan Plateau. Elasticity values of each vital rate accumulated through all sizes are given in the tables in the upper right corner.

### Contributions of vital rates to changes in population growth rates under warming

To examine how differences in *λ* under ambient and warming conditions were caused by warming-induced changes in vital rates, we conducted the Life Table Response Experiment (LTRE). We found that the reduction in *λ* of *E. nutans* under warming was mainly caused by increased shrinkage of tall individuals and reduced positive growth of short individuals (Figure 4a). Despite the high survival elasticity in *E. nutans*, the reduction in survival made only a small contribution to the reduction in its *λ* value (Figure 4a). On the other hand, the increase in *λ* of *H. tibeticum* under warming was mainly achieved by increased survival of large individuals. Although shrinkage made a considerable negative contribution to its population maintenance, this was offset by the positive contribution from increased survival under warming (Figure 4b). Fecundity contributed the least to the variation in *λ* in both species (Figure 4a,b).

**Figure 4.**
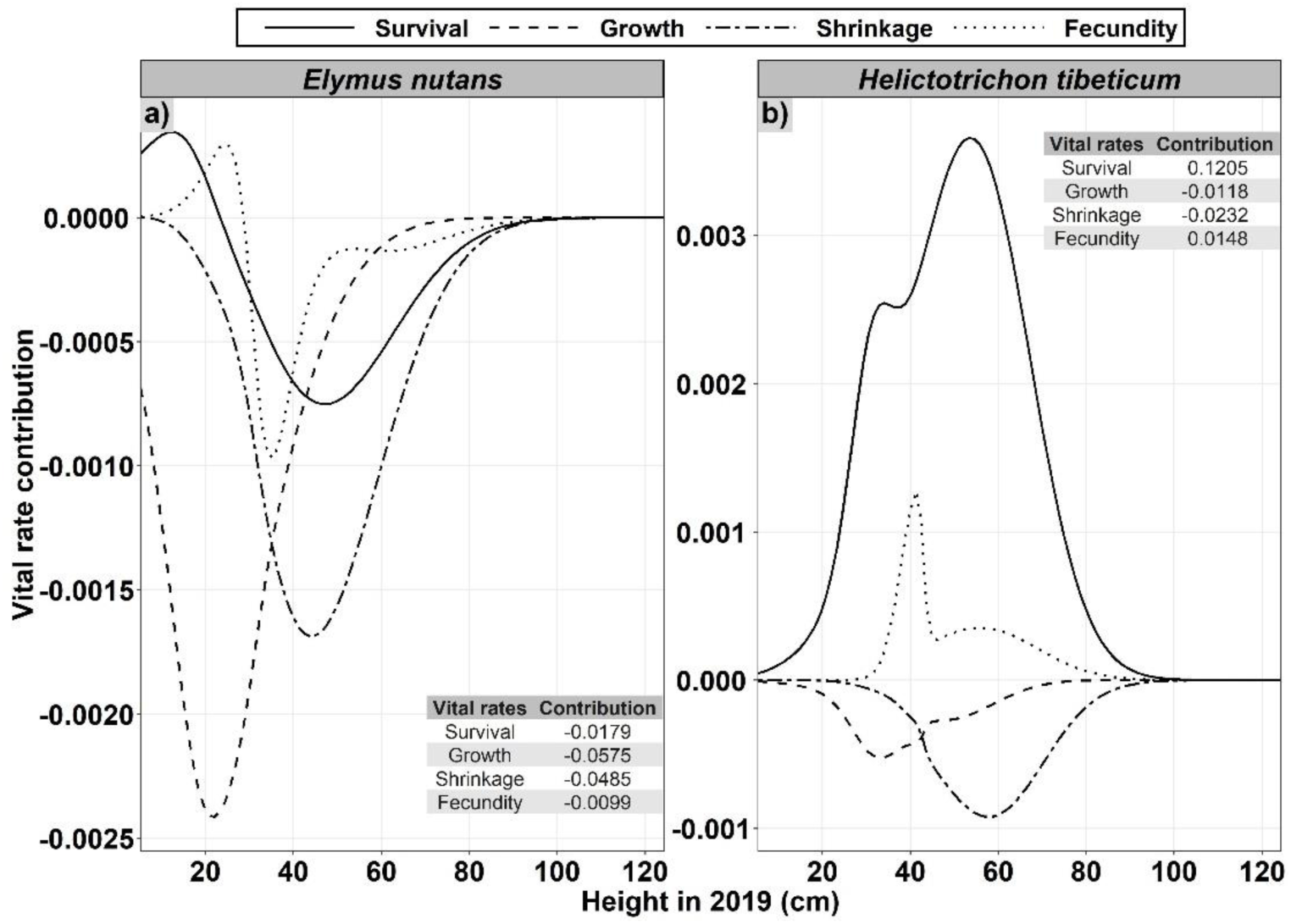
Results of Life Table Response Experiment analyses to examine the effects of a decade of warming treatment on the vital rate contribution of two co-occurring alpine herbaceous species, *Elymus nutans* (a) and *Helictotrichon tibeticum* (b), on the Tibetan Plateau. Cumulative contributions of each vital rate across all sizes to the warming-induced changes in population growth rates are given in the tables in the lower right corner (a) and upper right corner (b), respectively.

## DISCUSSION

### Population dynamics under climate warming

Quantitatively assessing how climate warming might affect the persistence of natural populations is critical for effective biodiversity conservation (Nomoto & Alexander, 2021; Sheth & Angert, 2018). Comparing the demographic responses of functionally similar species inhabiting the same habitats under elevated temperature can help evaluate the generality of the findings about the impacts of climate warming on population viability. However, this approach is rarely carried out (but see Nomoto & Alexander, 2021; Reed et al., 2021). Here, we examined the demographic responses of two co-occurring alpine herbaceous plants, *Elymus nutans* and *Helictotrichon tibeticum*, to a decade of *in situ* active warming on the Tibetan Plateau. Although the two species exhibited similar population growth rates under ambient conditions, *E. nutans* was projected to decline sharply and *H. tibeticum* to remain stable under warmed conditions. Different demographic responses of functionally similar species to environmental variability have been documented in diverse systems, including herbaceous plants in moist meadows (Jongejans & de Kroon, 2005), forb species in grasslands (Olsen et al., 2016), bunchgrass species in prairies (DeMarche et al., 2021), and congeneric endangered grassland herbs (Münzbergová, 2013). Such different responses varied not only in terms of magnitude, but even in direction (DeMarche et al., 2021; Jongejans & de Kroon, 2005; Münzbergová, 2013; Olsen et al., 2016). These findings, together with our results, suggest that coexisting species may face contrasting fates under climate change, resulting in community re-assembly, as it has been found in recent studies (Adler et al., 2012; Cant et al., 2021). Therefore, we may not easily transfer the demographic information obtained from one species to another inhabiting the same habitat, even those from the same functional group.

### Demographic processes drive population dynamics under climate warming

Climate warming may affect population viability by unevenly affecting different demographic processes. Demographic compensation among vital rates may allow species to maintain viable populations under changing environments (Doak & Morris, 2010; Peterson et al., 2018). In our study, both species exhibited decreased individual growth and increased shrinkage after a decade of warming, suggesting that the growing conditions became less favorable. However, the population of *H. tibeticum* successfully compensated for the negative effects of warming on growth by increased survival and reproduction, resulting in an increase in its overall population growth rate. Contrastingly, the population of *E. nutans* failed compensate, and was projected to decline sharply under warmed conditions. Demographic compensation at elevated temperatures has also been reported in a recent study on a perennial herb (Sheth & Angert, 2018). In that study, the authors found that adequate demographic compensation among vital rates enabled *Erythranthe cardinalis* to maintain viable populations in high latitude areas, while populations with insufficient compensation at low latitudes were projected to decline as climate change progresses. Together, these findings and our results suggest that demographic compensation may help populations to buffer warming induced negative effects and better adapt to warmed living conditions. Species lacking demographic compensation might be more likely to face future population declines.

The influence of changes in vital rates on overall population performance depends on the sensitivity of the population growth rate to such changes (de Kroon et al., 2000). In our study, adult survival in both species had the overwhelmingly highest elasticity values under both ambient and warmed conditions, suggesting that populations are especially sensitive to changes in adult survival. Indeed, our LTRE results showed that the contrasting fates of the two co-occurring species under warming were mainly attributable to the different responses of adult survival to warming. Therefore, our results suggest that increased mortality could lead to detrimental effects on population viability. Thus, more attention should be paid to adult survival in the face of climate change (McDonald et al., 2017; Sheth & Angert, 2018), something that may be difficult to achieve in short-term studies.

Climate warming may affect the distribution of elasticity values over a life cycle. In our study, the highest elasticity values of survival shifted toward shorter individuals under warming, indicating that maintaining the survival of small plants became more important for population persistence in the face of climate warming. Previous studies have reported that low winter temperatures in alpine regions could impose a strong selective pressure on plant overwintering, especially on seedlings and small individuals with less storage of carbon resources (Zait et al., 2020). Here, shorter individuals had much lower survival than taller ones, which might be partly due to the low capacity of small individuals to overwinter (Simons et al., 2010). Warming promoted the survival of shorter individuals in both species, indicating that future climate warming may benefit these small plant individuals by improving their capacity to overwinter.

The demographic responses to climate warming may be regulated by lifespan, a key life history trait that may affect a species’ capacity to cope with environmental variation (Garcia et al., 2008; Morris et al., 2008; Paniw et al., 2018). Short-lived species are often more sensitive to climate variability than long-lived species (Morris et al., 2018; Paniw et al., 2018). This pattern may be driven by short-lived species’ higher temporal variation in annual population growth rate, which may negatively affect the long-term stochastic population growth rate (Morris et al., 2008; Pfister, 1998). In our study, warming reduced the population growth of *E. nutans* (with a shorter life expectancy), but promoted the population growth of *H. tibeticum* (with a longer life expectancy). Recently, based on demographic data from 62 plant species in temperate biomes, Compagnoni et al. (2021) found that warming had stronger negative effects on the population growth rate of species with shorter generation time. Additionally, similar findings were also reported in birds and amphibians; short-lived species tended to show more pronounced responses to climate change, and species with a fast life history could be more vulnerable than species with a slower life history (Cayuela et al., 2017; Couet et al., 2022). Our results, together with these similar findings across diverse systems, suggest that lifespan might be key predictor of species’ susceptibility to climate change.

### Management implications in the face of future climate change

Livestock grazing is a common land use activity in alpine grassland ecosystems (Zhang et al., 2023). The grasslands on the Tibetan Plateau serve as an important resource for yaks (Long et al., 2008). We found that survival was key for population maintenance of both species, suggesting that grazing activities should be controlled at a level that minimizes mortality. Additionally, increased shrinkage under warming made a considerable negative contribution to the population growth rates of both species in the current study. Therefore, grazing activities should also be managed to a certain extent to avoid large increases in shrinkage. As survival and shrinkage are relatively easy to monitor, they could potentially serve as warning signals for the management of alpine grasslands under climate change.

Management strategies should be tailored to the species level as much as possible. Our study site was grazed annually in winter to mimic historic grazing regimes in the local region. The populations of *E. nutans* declined under both ambient and warmed conditions, implying that the current grazing regime is unsustainable in either current or future climates. Adjustments to the grazing regime may help to reverse population decline (Andrello et al., 2012). Further studies are needed to determine the optimal grazing regime, in terms of intensity and season, to maintain viable populations for *E. nutans* in face of climate change. Contrastingly, our findings on *H. tibeticum* suggest that the current grazing regime is likely sustainable for this species under future climate warming. However, the current study only examined the effects of 2℃ warming, the potential effects of further warming remain unknown.

Population management under climate change needs to account for the potential demographic impacts of extreme weather events. As observed in a study by Morris and Doak (2010), demographic rates may be improved in moderately warmer years but decline in the warmest years. Therefore, increased climate warming exceeding a tipping point may still be detrimental and drive future population declines (Morris & Doak, 2010; Peterson et al., 2018). Similarly, Keller and Shea (2021) found nonlinear effects of warming on different life history stages. Furthermore, Andrello et al. (2012) found that a heatwave reduced the population growth of a vulnerable perennial species, *Eryngium alpinum*, under all grazing regimes, and particularly exacerbated extinction risk under spring grazing. Therefore, the management of alpine plant populations in face of climate change also needs to take extreme weather events into account and adjust grazing activities accordingly.

Communities at high latitudes and at high elevations, like the Tibetan plateau system studied here, serve as bellwethers for future climates. Long-term warming experiments in such systems offer lessons for ecologists, land managers and conservationists keen to predict the effects of climate warming on plant populations’ long-term viability. We show that two similar, widely distributed alpine grasses had disparate responses to elevated temperature. This finding highlights the need for species-level investigation, particularly for economically important species such as these valuable forage plants.

## ACKNOWLEDGEMENTS

We acknowledge funding from the National Science Foundation of China (31971423).

## AUTHORS’ CONTRIBUTIONS

S-LL RSG, KS and J-SH conceived the study. H-TM collected the data, constructed models, wrote first draft of the manuscript, and all authors contributed substantially to revisions. All authors approved the final manuscript.

## DATA ACCESSIBILITY STATEMENT

All data used in this manuscript will be archived in Dryad and the data DOI will be provided should the manuscript be accepted for publication.

## REFERENCES

Adler, P. B., Dalgleish, H. J. & Ellner, S. P. (2012). Forecasting plant community impacts of climate variability and change: when do competitive interactions matter? Journal of Ecology, 100, 478–487. 10.1111/j.1365-2745.2011.01930.x

Andrello, M., Bizoux, J. P., Barbet-Massin, M., Gaudeul, M., Nicole, F. & Till-Bottraud, I. (2012). Effects of management regimes and extreme climatic events on plant population viability in *Eryngium alpinum*. Biological Conservation, 147, 99–106. 10.1016/j.biocon.2011.12.012

Anderson, J. T. & Wadgymar, S. M. (2020). Climate change disrupts local adaptation and favours upslope migration. Ecology Letters, 23, 181–192. 10.1111/ele.13427

Blumenthal, D. M., Kray, J. A., Ortmans, W., Ziska, L. H. & Pendall, E. (2016). Cheatgrass is favored by warming but not CO_2_ enrichment in a semi-arid grassland. Global Change Biology, 22, 3026–3038. 10.1111/gcb.13278

Cant, J., Salguero-Gómez, R., Kim, S. W., Sims, C. A., Sommer, B., Brooks, M., Malcolm, H. A., Pandolfi, J. M. & Beger, M. (2021). The projected degradation of subtropical coral assemblages by recurrent thermal stress. Journal of Animal Ecology, 90, 233–247. 10.1111/1365-2656.13340

Cayuela, H., Joly, P., Schmidt, B. R., Pichenot, J., Bonnaire, E., Priol, P., Peyronel, O., Laville, M. & Besnard, A. (2017). Life history tactics shape amphibians’ demographic responses to the North Atlantic Oscillation. Global Change Biology, 23, 4620–4638. 10.1111/gcb.13672

Caswell, H. (1989). Analysis of life table response experiments: Decomposition of effects on population growth rate. Ecological Modelling, 46, 221–237. 10.1016/0304-3800(89)90019-7

Caswell, H. (2001). Matrix Population Models: Construction, Analysis, and Interpretation (2nd ed.). Sunderland, MA: Sinauer Associates Inc.

Chen, H., Zhu, Q. A., Peng, C. H., Wu, N., Wang, Y. F., Fang, X. Q., Gao, Y. H., Zhu, D., Yang, G., Tian, J. Q., Kang, X. M., Piao, S. L., Ouyang, H., Xiang, W. H., Luo, Z. B., Jiang, H., Song, X. Z., Zhang, Y., Yu, G. R., … Wu, J. H. (2013). The impacts of climate change and human activities on biogeochemical cycles on the Qinghai-Tibetan Plateau. Global Change Biology, 19, 2940–2955. 10.1111/gcb.12277

Compagnoni, A., Levin, S., Childs, D.Z., Harpole, S., Paniw, M., Romer, G., Burns, J. H., Che-Castaldo, J., Ruger, N., Kunstler, G., Bennett, J. M., Archer, C. R., Jones, O. R., Salguero-Gómez, R. & Knight, T. M. (2021). Herbaceous perennial plants with short generation time have stronger responses to climate anomalies than those with longer generation time. Nature Communications, 12, 1824. 10.1038/s41467-021-21977-9

Couet, J., Marjakangas, E. L., Santangeli, A., Kalas, J. A., Lindstrom, A. & Lehikoinen, A. (2022). Short-lived species move uphill faster under climate change. Oecologia, 198, 877–888. 10.1007/s00442-021-05094-4

Cribari-Neto, F. & Zeileis, A. (2010). Beta Regression in R. Journal of Statistical Software, 34, 1–24. 10.18637/jss.v034.i02

Cui, H. J., Topper, J. P., Yang, Y., Vandvik, V. & Wang, G. X. (2018). Plastic Population Effects and Conservative Leaf Traits in a Reciprocal Transplant Experiment Simulating Climate Warming in the Himalayas. Frontiers in Plant Science, 9, 1069. 10.3389/fpls.2018.01069

de Kroon, H., van Groenendael, J. & Ehrlen, J. (2000). Elasticities: A review of methods and model limitations. Ecology, 81, 607–618. 10.1890/0012-9658(2000)081[0607:EAROMA]2.0.CO;2

DeMarche, M. L., Bailes, G., Hendricks, L. B., Pfeifer-Meister, L., Reed, P. B., Bridgham, S. D., Johnson, B. R., Shriver, R., Waddle, E., Wroton, H., Doak, D. F., Roy, B. A. & Morris, W. F. (2021). Latitudinal gradients in population growth do not reflect demographic responses to climate. Ecological Applications, 31, e02242. 10.1002/eap.2242

DeMarche, M. L., Doak, D. F. & Morris, W. F. (2018). Both life-history plasticity and local adaptation will shape range-wide responses to climate warming in the tundra plant *Silene acaulis*. Global Change Biology, 24, 1614–1625. 10.1111/gcb.13990

Doak, D. F. & Morris, W. F. (2010). Demographic compensation and tipping points in climate-induced range shifts. Nature, 467, 959–962. 10.1038/nature09439

Easterling, M. R., Ellner, S. P. & Dixon, P. M. (2000). Size-specific sensitivity: Applying a new structured population model. Ecology, 81, 694–708. 10.1890/0012-9658(2000)081[0694:SSSAAN]2.0.CO;2

Ellner, S. P., Childs, D. Z. & Rees, M. (2016). Data-driven Modelling of Structured Populations: A Practical Guide to the Integral Projection Model. Springer International Publishing, Switzerland.

Efron, B. (1987). Better bootstrap confidence-intervals. Journal of the American Statistical Association, 82, 171–185. 10.2307/2289144

Evers, S. M., Knight, T. M., Inouye, D. W., Miller, T. E. X., Salguero-Gómez, R., Iler, A. M. & Compagnoni, A. (2021). Lagged and dormant season climate better predict plant vital rates than climate during the growing season. Global Change Biology, 27, 1927–1941. 10.1111/gcb.15519

Fox J. (2016). Applied Regression Analysis and Generalized Linear Models (3rd ed.). Sage Publications, Inc.

Fuller, R. J., Williamson, T., Barnes, G. & Dolman, P. M. (2017). Human activities and biodiversity opportunities in pre-industrial cultural landscapes: relevance to conservation. Journal of Applied Ecology, 54, 459–469. 10.1111/1365-2664.12762

Garcia, M. B., Pico, F. X. & Ehrlen, J. (2008). Life span correlates with population dynamics in perennial herbaceous plants. American Journal of Botany, 95, 258–262. 10.3732/ajb.95.2.258

Gornish, E. S. (2014). Demographic effects of warming, elevated soil nitrogen and thinning on the colonization of a perennial plant. Population Ecology, 56, 645–656. 10.1007/s10144-014-0442-5

Griffith, A. B. (2017). Perturbation approaches for integral projection models. Oikos, 126, 1675–1686. 10.1111/oik.04458

Iler, A. M., CaraDonna, P. J., Forrest, J. R. K. & Post, E. (2021). Demographic Consequences of Phenological Shifts in Response to Climate Change. Annual Review of Ecology, Evolution, and Systematics, 52, 221–245. 10.1146/annurev-ecolsys-011921-032939

Jones, O. R., Barks, P., Stott, I., James, T. D., Levin, S., Petry, W. K., Capdevila, P., Che-Castaldo, J., Jackson, J., Romer, G., Schuette, C., Thomas, C. C. & Salguero-Gómez, R. (2022). Rcompadre and Rage-Two R packages to facilitate the use of the COMPADRE and COMADRE databases and calculation of life-history traits from matrix population models. Methods in Ecology and Evolution, 13, 770–781. 10.1111/2041-210X.13792

Jongejans, E. & De Kroon, H. (2005). Space versus time variation in the population dynamics of three co-occurring perennial herbs. Journal of Ecology, 93, 681–692. 10.1111/j.1365-2745.2005.01003.x

Keller, J. A. & Shea, K. (2021). Warming and shifting phenology accelerate an invasive plant life cycle. Ecology, 102, e03219. 10.1002/ecy.3219

Li, S. L., Yu, F. H., Werger, M. J. A., Dong, M. & Zuidema, P. A. (2011). Habitat-specific demography across dune fixation stages in a semi-arid sandland: understanding the expansion, stabilization and decline of a dominant shrub. Journal of Ecology, 99, 610–620. 10.1111/j.1365-2745.2010.01777.x

Liu, H. Y., Mi, Z. R., Lin, L., Wang, Y. H., Zhang, Z. H., Zhang, F. W., Wang, H., Liu, L. L., Zhu, B. A., Cao, G. M., Zhao, X. Q., Sanders, N. J., Classen, A. T., Reich, P. B. & He, J. S. (2018). Shifting plant species composition in response to climate change stabilizes grassland primary production. Proceedings of the National Academy of Sciences, 115, 4051–4056. 10.1073/pnas.1700299114

Long, R. J., Ding, L. M., Shang, Z. H. & Guo, X. H. (2008). The yak grazing system on the Qinghai-Tibetan plateau and its status. Rangeland Journal, 30, 241–246. 10.1071/RJ08012

Lucas, R. W., Forseth, I. N. & Casper, B. B. (2008). Using rainout shelters to evaluate climate change effects on the demography of *Cryptantha flava*. Journal of Ecology, 96, 514–522. 10.1111/j.1365-2745.2007.01350.x

McDonald, J. L., Franco, M., Townley, S., Ezard, T. H. G., Jelbert, K. & Hodgson, D. J. (2017). Divergent demographic strategies of plants in variable environments. Nature ecology & evolution, 1, 0029. 10.1038/s41559-016-0029

McLean, N., Lawson, C. R., Leech, D. I. & van de Pol, M. (2016). Predicting when climate-driven phenotypic change affects population dynamics. Ecology Letters, 19, 595–608. 10.1111/ele.12599

IPCC. (2019). Polar Regions. In: IPCC Special Report on the Ocean and Cryosphere in a Changing Climate (Pörtner, H. O., Roberts, D. C., Masson-Delmotte, V., Zhai, P., M. Tignor, M., Poloczanska, M. E., Mintenbeck, K., Alegría, A., Nicolai, M., Okem, A., Petzold, J., Rama, B. & Weyer, N. M., Eds.). Cambridge University Press. 10.1017/9781009157964

Morris, W. F., Pfister, C. A., Tuljapurkar, S., Haridas, C. V., Boggs, C. L., Boyce, M. S., Bruna, E. M., Church, D. R., Coulson, T., Doak, D. F., Forsyth, S., Gaillard, J. M., Horvitz, C. C., Kalisz, S., Kendall, B. E., Knight, T. M., Lee, C. T. & Menges, E. S. (2008). Longevity can buffer plant and animal populations against changing climatic variability. Ecology, 89, 19–25. 10.1890/07-0774.1

Münzbergová, Z. (2013). Comparative demography of two co-occurring *Linum* species with different distribution patterns. Plant Biology, 15, 963–970. 10.1111/plb.12007

Nomoto, H. A. & Alexander, J. M. (2021). Drivers of local extinction risk in alpine plants under warming climate. Ecology Letters, 24, 1157–1166. 10.1111/ele.13727

Olsen, S. L., Topper, J. P., Skarpaas, O., Vandvik, V. & Klanderud, K. (2016). From facilitation to competition: temperature-driven shift in dominant plant interactions affects population dynamics in seminatural grasslands. Global Change Biology, 22, 1915–1926. 10.1111/gcb.13241

Panetta, A. M., Stanton, M. L. & Harte, J. (2018). Climate warming drives local extinction: Evidence from observation and experimentation. Science Advance, 4, eaaq1819. 10.1126/sciadv.aaq1819

Paniw, M., Ozgul, A. & Salguero-Gómez, R. (2018). Interactive life-history traits predict sensitivity of plants and animals to temporal autocorrelation. Ecology Letters, 21, 275–286. 10.1111/ele.12892

Pearson, R. G., Stanton, J. C., Shoemaker, K. T., Aiello-Lammens, M. E., Ersts, P. J., Horning, N., Fordham, D. A., Raxworthy, C. J., Ryu, H. Y., McNees, J. & Akcakaya, H. R. (2014). Life history and spatial traits predict extinction risk due to climate change. Nature Climate Change, 4, 217–221. 10.1038/nclimate2113

Peterson, M. L., Doak, D. F. & Morris, W. F. (2018). Both life-history plasticity and local adaptation will shape range-wide responses to climate warming in the tundra plant *Silene acaulis*. Global Change Biology, 24, 1614–1625. 10.1111/gcb.13990

Pfister, C. A. (1998). Patterns of variance in stage-structured populations: evolutionary predictions and ecological implications. Proceedings of the National Academy of Sciences of the United States of America, 95, 213–218. 10.1073/pnas.95.1.213

R Core Team. (2021) R: A language and environment for Statistical Computing. R Foundation for Statistical Computing. https://www.r-project.org/

Reed, P. B., Peterson, M. L., Pfeifer-Meister, L. E., Morris, W. F., Doak, D. F., Roy, B. A., Johnson, B. R., Bailes, G. T., Nelson, A. A. & Bridgham, S. D. (2021). Climate manipulations differentially affect plant population dynamics within versus beyond northern range limits. Journal of Ecology, 109, 664–675. 10.1111/1365-2745.13494

Salguero-Gómez, R. & Casper, B. B. (2010). Keeping plant shrinkage in the demographic loop. Journal of Ecology, 98, 312–323. 10.1111/j.1365-2745.2009.01616.x

Selwood, K. E., McGeoch, M. A. & Nally, R. (2015). The effects of climate change and land-use change on demographic rates and population viability. Biological Reviews, 90, 837–853. 10.1111/brv.12136

Sheth, S. N. & Angert, A. L. (2018). Demographic compensation does not rescue populations at a trailing range edge. Proceedings of the National Academy of Sciences of the United States of America, 115, 2413–2418. 10.1073/pnas.1715899115

Simons, A. M, Goulet, J. M. & Bellehumeur, K. F. (2010). The effect of snow depth on overwinter survival in *Lobelia inflata*. Oikos, 119, 1685–1689. 10.1111/j.1600-0706.2010.18515.x

Struckman, S., Couture, J. J., LaMar, M. D. & Dalgleish, H. J. (2019). The demographic effects of functional traits: an integral projection model approach reveals population-level consequences of reproduction-defence trade-offs. Ecology Letters, 22, 1396–1406. 10.1111/ele.13325

Tenhumberg, B., Crone, E. E., Ramula, S. & Tyre, A. J. (2018). Time-lagged effects of weather on plant demography: drought and *Astragalus scaphoides*. Ecology, 99, 915–925. 10.1002/ecy.2163

Verrall, B. & Pickering, C. M. (2020). Alpine vegetation in the context of climate change: A global review of past research and future directions. Science of the Total Environment, 748, 141344. 10.1016/j.scitotenv.2020.141344

Villellas, J., Doak, D. F., Garcia, M. B. & Morris, W. F. (2015). Demographic compensation among populations: what is it, how does it arise and what are its implications? Ecology Letters, 18, 1139–1152. 10.1111/ele.12505

Wang, H., Liu, H., Cao, G., Ma, Z., Li, Y., Zhang, F., Zhao, X., Zhao, X., Jiang, L., Sanders, N. J., Classen, A. T. & He, J. S. (2020). Alpine grassland plants grow earlier and faster but biomass remains unchanged over 35 years of climate change. Ecology Letters, 23, 701–710. 10.1111/ele.13474

Wang, Y., Lv, W., Xue, K., Wang, S., Zhang, L., Hu, R., Zeng, H., Xu, X., Li, Y., Jiang, L., Hao, Y., Du, J., Sun, J., Dorji, T., Piao, S., Wang, C., Luo, C., Zhang, Z., Chang, X., … Niu, H. (2022). Grassland changes and adaptive management on the Qinghai-Tibetan Plateau. Nature Reviews Earth & Environment, 3, 668–683. 10.1038/s43017-022-00330-8

Wilcoxon, F. (1945). Individual comparisons by ranking methods. Biometrics, 1, 80–83. 10.2307/3001968

Zait, Y., Konsens, I. & Schwartz, A. (2020). Elucidating the limiting factors for regeneration and successful establishment of the thermophilic tree *Ziziphus spina-christi* under a changing climate. Scientific Reports, 10, 14335. 10.1038/s41598-020-71276-4

Zhang, R., Gallagher, R. S. & Shea, K. (2012). Maternal warming affects early life stages of an invasive thistle. Population Biology, 14, 738–788. 10.1111/j.1438-8677.2011.00561.x

Zhang, Z., Zhao, Y., Lin, H., Li, Y., Fu, J., Wang, Y., Sun, J. & Zhao, Y. (2023). Comprehensive analysis of grazing intensity impacts alpine grasslands across the Qinghai-Tibetan Plateau: A meta-analysis. Frontiers in Plant Science, 13, 1083709. 10.3389/fpls.2022.1083709

Zuidema, P. A. & Franco, M. (2001). Integrating vital rate variability into perturbation analysis: an evaluation for matrix population models of six plant species. Journal of Ecology, 89, 995–1005. 10.1111/j.1365-2745.2001.00621.x

