## Supplemental File for "Differences in adult survival drive divergent demographic responses to a decade of field warming on the Tibetan Plateau"

**Supporting information**

**Table S1** The fixed effects of warming treatment and initial plant height on the vital rates of two co-occurring alpine herbaceous species, *Elymus nutans* and *Helictotrichon tibeticum*, on the Tibetan Plateau. Wald chi-square tests are provided for the fixed effects in each vital rate model and the significance was determined using type-II tests.

| **Species** | **Vital rates** | **Parameters** | ***Wald Chi-Square*** | | ***P*** |
| --- | --- | --- | --- | --- | --- |
| *E. nutans* | Survival probability | Plant height | 60.34 | <0.01 | |
|  |  | Treatment | 329.22 | < 0.01 | |
|  |  | Plant height × treatment | 298.60 | < 0.01 | |
|  | Changes in height | Plant height | 33535.26 | < 0.01 | |
|  |  | Treatment | 40587.46 | < 0.01 | |
|  |  | Plant height × treatment | 2113.71 | < 0.01 | |
|  | Probability of reproduction | Plant height | 25596.63 | < 0.01 | |
|  |  | Treatment | 553.40 | < 0.01 | |
|  |  | Plant height × treatment | 1945.42 | < 0.01 | |
|  | Number of seeds | Plant height | 6128.65 | < 0.01 | |
|  |  | Treatment | 1652.39 | < 0.01 | |
|  |  | Plant height × treatment | 4954.75 | < 0.01 | |
| *H. tibeticum* | Survival probability | Plant height | 866.36 | < 0.01 | |
|  |  | Treatment | 3471.59 | < 0.01 | |
|  |  | Plant height × treatment | 50.30 | < 0.01 | |
|  | Changes in height | Plant height | 33235.05 | < 0.01 | |
|  |  | Treatment | 11806.93 | < 0.01 | |
|  |  | Plant height × treatment | 58.67 | < 0.01 | |
|  | Probability of reproduction | Plant height | 4218.85 | < 0.01 | |
|  |  | Treatment | 1135.74 | < 0.01 | |
|  |  | Plant height × treatment | 59.22 | < 0.01 | |
|  | Number of spikes | Plant height | 2681.84 | < 0.01 | |
|  |  | Treatment | 2733.51 | < 0.01 | |
|  |  | Plant height × treatment | 23.34 | < 0.01 | |

**Table S2** The effects of warming treatment on the average plant height, crown area and life expectancy with 95% confident intervals for two co-occurring alpine herbaceous species, *Elymus nutans* and *Helictotrichon tibeticum*, on the Tibetan Plateau. Values in brackets refer to 95% confidence intervals (CI).

| **Species** | **Treatment** | **Height (cm)** | **Crown area (cm^2^)** | **Life expectancy (years)** |
| --- | --- | --- | --- | --- |
| *E. nutans* | Ambient | 23.70 [19.88, 27.52] | 22.94 [15.39, 30.48] | 4.47 [4.42, 4.53] |
|  | Warming | 22.77 [19.40, 26.14] | 24.30 [17.38, 31.22] | 2.99 [2.96, 3.01] |
| *H. tibeticum* | ambient | 33.3 [29.66, 37.06] | 112.10 [72.64, 151.57] | 6.68 [6.52, 6.76] |
|  | Warming | 34.56 [29.52, 39.60] | 80.74 [46.12, 115.37] | 15.43 [15.11, 16.01] |

**Table S3** The effects of warming treatment on the occurrence and intensity in plant height of two co-occurring alpine herbaceous species, *Elymus nutans* and *Helictotrichon tibeticum*, on the Tibetan Plateau. We modeled the proportion of shrinkage occurrence using a generalized linear model from the beta family with a logit-link function, and warming treatment was included as an independent variable. We applied a Wald chi-square test to test the significance of warming treatment in the model. We performed a Wilcoxon signed rank test to estimate the warming effect on intensity of shrinkage.

| **Species** |  | **Treatment** | **Mean value** | ***χ^2^*** | ***W*** | ***P*** |
| --- | --- | --- | --- | --- | --- | --- |
| *E. nutans* | Proportional occurrence of shrinkage (%) | Ambient | 26.69 | 5.630 |  | 0.018 |
|  |  | Warming | 38.65 |  |  |  |
|  | Intensity of shrinkage (cm) | Ambient | -17.26 |  | 1230 | 0.814 |
|  |  | Warming | -18.67 |  |  |  |
| *H. tibeticum* | Proportional occurrence of shrinkage (%) | Ambient | 16.24 | 2.602 |  | 0.107 |
|  |  | Warming | 23.95 |  |  |  |
|  | Intensity of shrinkage (cm) | Ambient | -13.36 |  | 1132 | 0.874 |
|  |  | Warming | -14.51 |  |  |  |

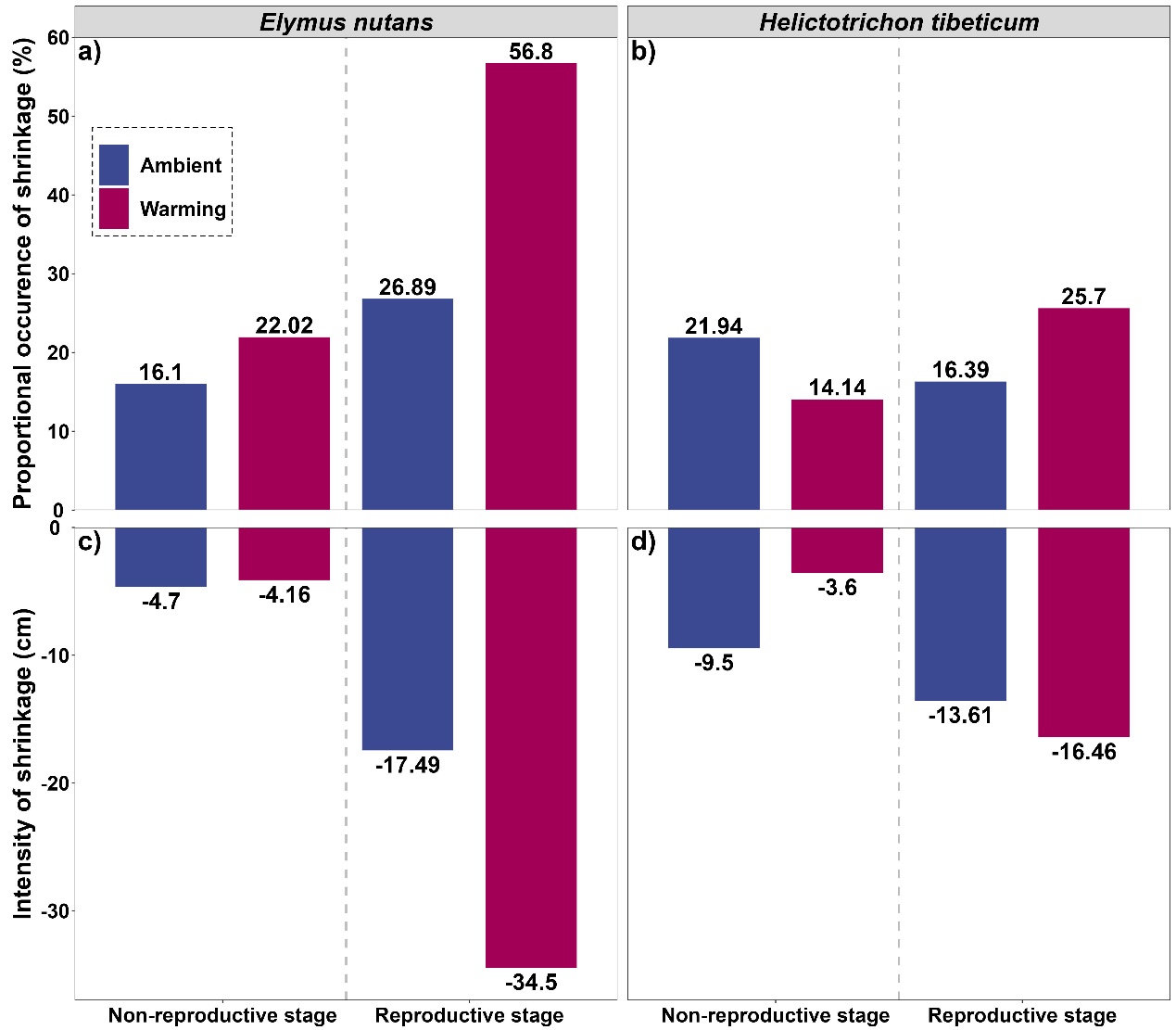

**Figure S1** The effects of a decade of warming treatment on the occurrence and intensity of shrinkage in plant height of two co-occurring alpine herbaceous species, *Elymus nutans* (a, c) and *Helictotrichon tibeticum* (b, d), on the Tibetan Plateau.
